# Genetics of recombination rate variation in the pig

**DOI:** 10.1101/2020.03.17.995969

**Authors:** Martin Johnsson, Andrew Whalen, Roger Ros-Freixedes, Gregor Gorjanc, Ching-Yi Chen, William O. Herring, Dirk-Jan de Koning, John M. Hickey

## Abstract

**Background:** In this paper, we estimated recombination rate variation within the genome and between individuals in the pig using multiocus iterative peeling for 150,000 pigs across nine genotyped pedigrees. We used this to estimate the heritability of recombination and perform a genome-wide association study of recombination in the pig.

**Results:** Our results confirmed known features of the pig recombination landscape, including differences in chromosome length, and marked sex differences. The recombination landscape was repeatable between lines, but at the same time, the lines also showed differences in average genome-wide recombination rate. The heritability of genome-wide recombination was low but non-zero (on average 0.07 for females and 0.05 for males). We found three genomic regions associated with recombination rate, one of them harbouring the *RNF212* gene, previously associated with recombination rate in several other species.

**Conclusion:** Our results from the pig agree with the picture of recombination rate variation in vertebrates, with low but nonzero heritability, and a major locus that is homologous to one detected in several other species. This work also highlights the utility of using large-scale livestock data to understand biological processes.

## Background

This paper shows that recombination rate in the pig (*Sus scrofa*) is genetically variable and associated with alleles at the *RNF212* gene.

Recombination causes exchange of genetic material between homologous chromosomes. At meiosis, after chromosomes have been paired up and duplicated, they break and exchange pieces of chromosome arms. These recombinations are not evenly distributed along chromosomes. This gives rise to a variable recombination rate landscape with peaks and troughs.

The recombination rate landscape of the pig has been estimated previously [1]. It shows broadly the same features as in other mammals: low recombination rate in the centre of chromosomes, local hotspots of high recombination rate, a correlation between recombination rate and the fraction of guanine and cytosine bases (GC content), and sex difference in recombination rate [2–4]. In this paper, we investigated how recombination rate varied between individuals and populations in the pig.

Recombination rate is genetically variable in several other species. Studies in humans [5, 6], cattle [7–10], deer [11], sheep [12, 13] and chickens [14] have observed genetic influence on recombination rate, and genetic associations with alleles at a handful of genes involved in meiosis, including *RNF212*, *REC8* and *PRDM9* (reviewed by [15, 16]).

To be able to analyse the genetic basis of recombination, we need recombination estimates from a large number related pigs. Recombination rate can be estimated by phasing genotypes in pedigrees [17–20], by direct counting in gametes [21, 22], or by measuring linkage disequilibrium in population samples [2]. Counting methods require specific experiments to gather data. Linkage disequilibrium methods only provide averages for a population. In this paper, we used a new pedigree method based on multilocus iterative peeling [23, 24] to estimate recombination simultaneously with genotype imputation. This allowed us to use data from a pig breeding programme, where variable density genotype data has been gathered for genomic selection.

Our results confirmed known features of the pig recombination landscape, including differences in chromosome length, and marked sex difference. The recombination landscape was repeatable between lines, but at the same time, the lines showed differences in average genome-wide recombination rate. The heritability of genome-wide recombination was low but non-zero. We found three genomic regions associated with recombination rate, one of them harbouring the *RNF212* gene, previously associated with recombination rate in several other species.

## Methods

We estimated the recombination rate landscape in nine lines of pigs from a commercial breeding programme. We performed six analyses:

1. We estimated the average number of recombinations on each chromosome (the genetic length of chromosomes), and analysed between-sex and between-line differences in genetic length. We compared these estimates to previously published estimates.
2. We estimated the distribution of recombinations along chromosomes (recombination rate landscapes), and analysed between-line and between-sex differences.
3. We estimated the correlation between recombination rate and DNA sequence features previously known to correlate with recombination rate.
4. We estimated pedigree heritability and genomic heritability of recombination rate.
5. We ran a genome-wide association study to detect markers associated with recombination rate.
6. We ran a simulation to test the performance of the method.

### Data

We used SNP chip genotype data from nine lines of pigs from the Pig Improvement Company (PIC) breeding programme. This programme contains a diverse collection of genetics, which represent broadly used populations, including animals of Large White, Landrace, Duroc, Hampshire and Pietrain heritage. The pigs were genotyped at a mix of densities; either at low density (15K markers) using GGP-Porcine LD BeadChips (GeneSeek, Lincoln, NE) or at high density (60K or 75K markers) using GGP-Porcine HD BeadChips (GeneSeek, Lincoln, NE). In total, genotype data was available on 390,758 pigs.

### Recombination rate estimation using multilocus iterative peeling

We used multilocus iterative peeling to estimate the number and location of the recombination events in each individual [23, 24]. Multilocus iterative peeling uses pedigree data to calculate the phased genotype of each individual as a combination of information from the individual’s own genetic data, and that of their parents (anterior probabilities) and offspring (posterior probabilities) [25]. Multilocus iterative peeling builds on previous peeling algorithms by tracking which parental haplotype an individual inherits at each locus (segregation probabilities). This information can be used to determine which allele an individual inherits, particularly from parents who are heterozygous for that allele.

The segregation probabilities can be used to determine the number and location of likely recombination events. When a recombination happens, the offspring will inherit from a different parental haplotype. This will cause one, or both of the segregation probabilities to change, i.e. the segregation probability will change from a value close to 0 (likely to inherit the maternal haplotype) to 1 (likely to inherit the paternal haplotype). By analysing the joint distribution of neighbouring segregation probabilities, we are able to calculate the expected number of recombinations between two loci, and the expected number of recombinations across an entire chromosome.

To aid recombination rate estimation, we introduced two simplifications to the multilocus peeling method:

1. The segregation probabilities and the anterior probabilities were calculated separately for each parent in lieu of modelling their full joint distribution.
2. The segregation and genotype probabilities of the offspring were called when estimating the posterior term for each parent.

These simplifications were introduced to reduce runtime and memory requirements. In particular, by calling the segregation and genotype values, we are able to store many of the calculations in lookup tables instead of re-computing them for each locus, and each individual. In addition, the calling of segregation values reduced the chance that feedback loops occurred between offspring with fractional segregation values at multiple nearby loci.

A calling threshold of 0.99 was used for the segregation probabilities, and a calling threshold of 0.9 was used for the genotype probabilities. Segregation probabilities that did not reach the threshold were set to missing (equally likely to inherit either parental haplotype). Genotype probabilities that did not meet the threshold were also set to missing (all genotype states equally likely).

The joint distribution of segregation values depends on the chromosome length (in cM). To estimate chromosome length, we initialized the length to 100cM (on average 1 recombination per chromosome), and then refined this estimate in a series of steps. At each step we calculated the expected number of recombination for each individual at each locus, and set the chromosome length based on the average population recombination rate. This step was repeated four times. Preliminary simulations found that chromosome length estimates converged after four iterations, and that the recombination estimates for target individuals were insensitive to the assumed chromosome length.

### Filtering of individuals

After recombination estimation, we filtered the data by removing individuals without genotyped parents and grandparents in order to focus on those with high-quality recombination estimates. Filtering reduced the number of pigs to 145,763. Table 1 shows the resulting number of individuals per line post-filtering, and the total number of dams and sires for those individuals.

**Table 1.** Number of individuals that passed filtering in each line, and the unique number of their dams and sires. By necessity, we inferred recombination rates from an equal number of maternal and paternal chromosomes, but they derive from a much larger number of dams than sires.

| Line | Individuals kept | Dams | Sires |
| --- | --- | --- | --- |
| 1 | 23273 | 2651 | 437 |
| 2 | 16661 | 2255 | 368 |
| 3 | 14278 | 2169 | 215 |
| 4 | 7153 | 1239 | 163 |
| 5 | 33566 | 4349 | 293 |
| 6 | 11666 | 1971 | 162 |
| 7 | 263 | 76 | 20 |
| 8 | 4177 | 727 | 78 |
| 9 | 34726 | 5171 | 492 |

### Comparison between lines and to published maps

To compare the recombination landscapes of the nine lines we calculated pairwise correlations between lines of the estimated recombination rates at each marker interval, within each sex. To compare the recombination landscapes of the sexes, we calculated the correlation between sexes within each line.

We compared map length between lines using a linear model, fitting the number of recombinations observed on a chromosome as response, and fixed effects for each line and chromosome.

To compare the estimated landscapes to published landscapes, we also compared our results to the results of [1] by plotting our map length of each chromosome against published map lengths.

### Correlation with genome features

To investigate the relationship between local recombination rate and genomic features, we divided the autosomal part of the Sscrofa11.1 genome [26] into 2272 windows of 1 Mbp. We used Biostrings version 2.52.0 in the R statistical environment to estimate three features of sequence composition:

- fraction of guanine and cytosine bases (GC content);
- the PDRM9 consensus motif CCNCCNTNNCCNC [27];
- the CCCCACCCC motif, which was the most strongly associated with recombination in the pig in [1].

We used repeat data from RepeatMasker (http://www.repeatmasker.org) [28] from the pig genome to estimate the density of repeats in the same windows. We subdivided the total content of repeats into three broad categories:

- Fraction of LTR elements
- Fraction of DNA repeats elements
- Fraction of low complexity repeats

We calculated the correlation between the recombination rate and the sequence features within each window.

To find putative pericentromeric regions, we used the inferred centromere positions from [26]. On chromosomes 8, 11 and 15, where there were more than one inferred location far apart, we picked the most likely location based on karyotypes from [29].

### Heritability of genome-wide recombination rate

We estimated the narrow-sense heritability of genome-wide recombination rate using animal models in MCMCglmm [30] version 2.29. We estimated the heritability of recombination using genome-wide recombination rates per megabasepair. We fitted a pedigree animal model with an additive genetic effect and a permanent environmental effect for each parent as random effects. Because we measured recombination rate in parents of genotyped offspring, who have varying numbers of offspring (see Table 1), we used a model with repeated records and a permanent environmental effect for each parent. We analysed sexes and lines separately. We used parameter expanded priors [31] for the individual variance component and for the additive genetic variance component, using V = 1, ν = 1, α_μ_ = 0, α_V_ = 1000, which corresponds to a half-Cauchy prior with scale 100, and an inverse-Wishart prior (V = 1, ν = 1) for the residual variance. Because of the low number of dams and sires, we excluded the smallest line (line 7) from the quantitative genetic analysis. We also excluded parents with an extremely high average recombination rate (> 5 cM/Mbp).

### Genome-wide association

We performed genome-wide association studies of genome-wide recombination rates using hierarchical linear mixed models in RepeatABEL [32] version 1.1. The linear mixed model uses a genomic relationship matrix to account for relatedness while including a random permanent environmental effect for each parent. We analysed sexes and lines separately. We used imputed best-guess genotypes from the same run of AlphaPeel. Because of the low number of dams and sires, we again excluded line 7 from the analysis, and parents with average recombination rate > 5 cM/Mbp. We report significant markers below a conventional threshold of p < 5 · 10^−8^. We used the most significant marker in each region to report variance explained and the frequency of the allele associated with higher recombination. When there were more than one marker with the same p-value, we selected the marker closest to the middle of the interval.

### Simulations

To demonstrate that the method works, we tested it on a synthetic dataset with features similar to real data. We simulated genotype data with AlphaSimR 0.10.0. We simulated one chromosome, using the same pedigree and same number of genotyped markers as the largest line. The simulated recombination landscape had a constant recombination rate in the middle of the chromosome, and two regions of high recombination rate at the ends, described by second degree polynomials (the figure shows the resulting true recombination rate). We assessed accuracy of the inferred recombination landscape by calculating the correlation between the estimated number of recombination at each marker interval and the true number of recombination. We also calculated the correlation between the estimated number of recombinations and a smoothed recombination landscape, using a window of 50 markers.

## Results

Our results showed that:

1. There was variation in the genetic length of chromosomes between sexes and lines.
2. The recombination rate landscape was similar between lines but different between sexes.
3. We confirmed previous findings that local recombination rate is correlated with GC content, repeat content, the CCCCACCCC sequence motif, but not the previously described correlation with the PRDM9 consensus motif.
4. The heritability of recombination rate was on average 0.07 for females and 0.05 for males.
5. Three regions of the genome were associated with recombination rate, one of them containing the candidate gene *RNF212.*
6. In simulation, we found that multilocus iterative peeling could estimate the number of recombinations per individual with an accuracy of 0.7 for dams and 0.5 for sires, and the average recombination landscape along a chromosome, but with a tendency to overestimate the genetic length.

### Variation in genetic map length between lines and sexes

The genetic length of chromosomes was different between lines and sexes. Figure 1 shows the estimated map length of each chromosome, along with previously published estimates [1]. Table 2 gives the estimated of total map length in each sex and line, with confidence intervals derived from a linear model. On average, we estimated a sex-averaged map of 21.5 Morgan (0.95 cM/Mbp), a female map of 23.6 Morgan (1.04 cM/Mbp), and a male map of Morgan (0.86 cM/Mbp). Supplementary tables 1-3 contain male, female, and sex-averaged consensus maps of the pig recombination landscape.

**Figure 1.**
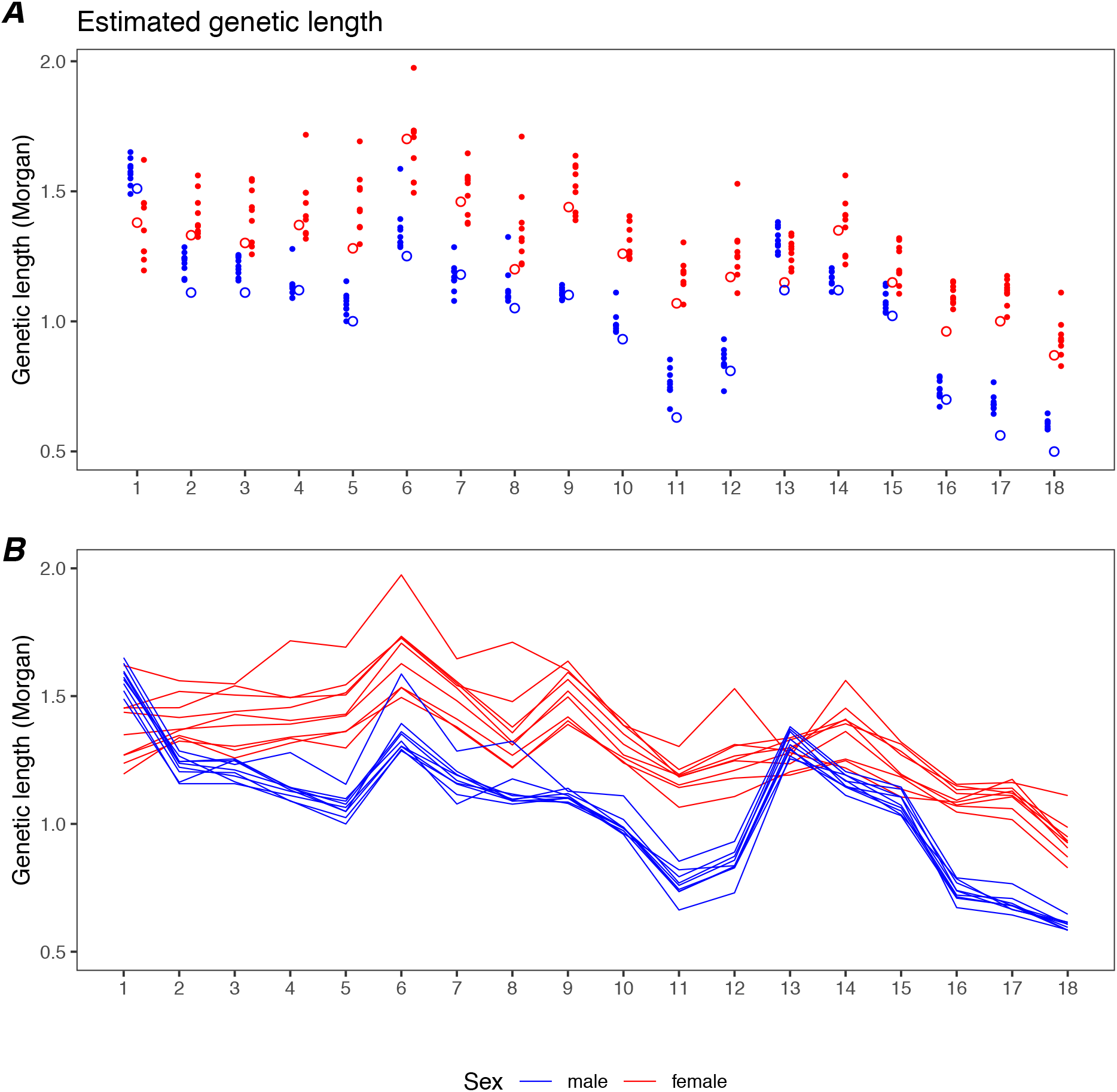
Genetic length of each pig autosome, as estimated by multilocus iterative peeling. The horizontal axis corresponds to chromosomes 1-18. Red dots and lines show female estimates, while blue dots and lines show male estimates. Panel A compares estimates from multilocus iterative peeling (filled dots) to estimates from [1] (open circles). Panel B shows estimates from the same line of pigs connected by lines.

**Table 2.** Estimates from linear model of total map length. Intervals are 95% confidence intervals.

| Line | Sex | Map length<br>(Morgan) | Lower | Upper | Rate<br>(cM/Mbp) |
| --- | --- | --- | --- | --- | --- |
| 1 | female | 23.6 | 23.5 | 23.6 | 1.04 |
| 1 | male | 19.4 | 19.4 | 19.5 | 0.86 |
| 2 | female | 24.1 | 24.1 | 24.2 | 1.06 |
| 2 | male | 20.0 | 20.0 | 20.0 | 0.88 |
| 3 | female | 22.3 | 22.2 | 22.3 | 0.98 |
| 3 | male | 18.2 | 18.1 | 18.2 | 0.80 |
| 4 | female | 23.5 | 23.4 | 23.5 | 1.04 |
| 4 | male | 19.3 | 19.3 | 19.4 | 0.85 |
| 5 | female | 22.8 | 22.7 | 22.8 | 1.01 |
| 5 | male | 18.7 | 18.6 | 18.7 | 0.82 |
| 6 | female | 23.7 | 23.6 | 23.7 | 1.04 |
| 6 | male | 19.5 | 19.5 | 19.6 | 0.86 |
| 7 | female | 25.9 | 25.5 | 26.2 | 1.14 |
| 7 | male | 21.7 | 21.4 | 22.1 | 0.96 |
| 8 | female | 24.1 | 24.0 | 24.2 | 1.06 |
| 8 | male | 20.0 | 19.9 | 20.1 | 0.88 |
| 9 | female | 22.6 | 22.6 | 22.6 | 1.00 |
| 9 | male | 18.5 | 18.4 | 18.5 | 0.82 |
| Average | female | 23.6 |  |  | 1.04 |
|  | male | 19.5 |  |  | 0.86 |
|  | sex-average | 21.5 |  |  | 0.95 |

Our estimated genetic lengths of chromosomes were comparable to previous estimates, but tended to be higher. We found that females have higher recombination rate, except on chromosome 1, where male recombination rate was higher, and chromosome 13, where the recombination rate is similar in both sexes. This confirms previous results [1].

### Difference in recombination landscape between sexes

The shape of the recombination landscape was similar between lines but different between sexes. Figure 2 presents the recombination rate landscape for each chromosome, and Figure 3 shows the correlation between the per-marker interval recombination rate estimates, between lines and between sexes. Both sexes had higher recombination rate near chromosome ends and lower recombination rate in the middle of the chromosomes. However, there were several broad regions of elevated female recombination rate which was not present in the males. These regions were repeatable between lines. The mean between-line correlation was 0.83 in females and 0.70 in males, whereas the mean correlation between sexes was 0.40 across lines.

**Figure 2.**
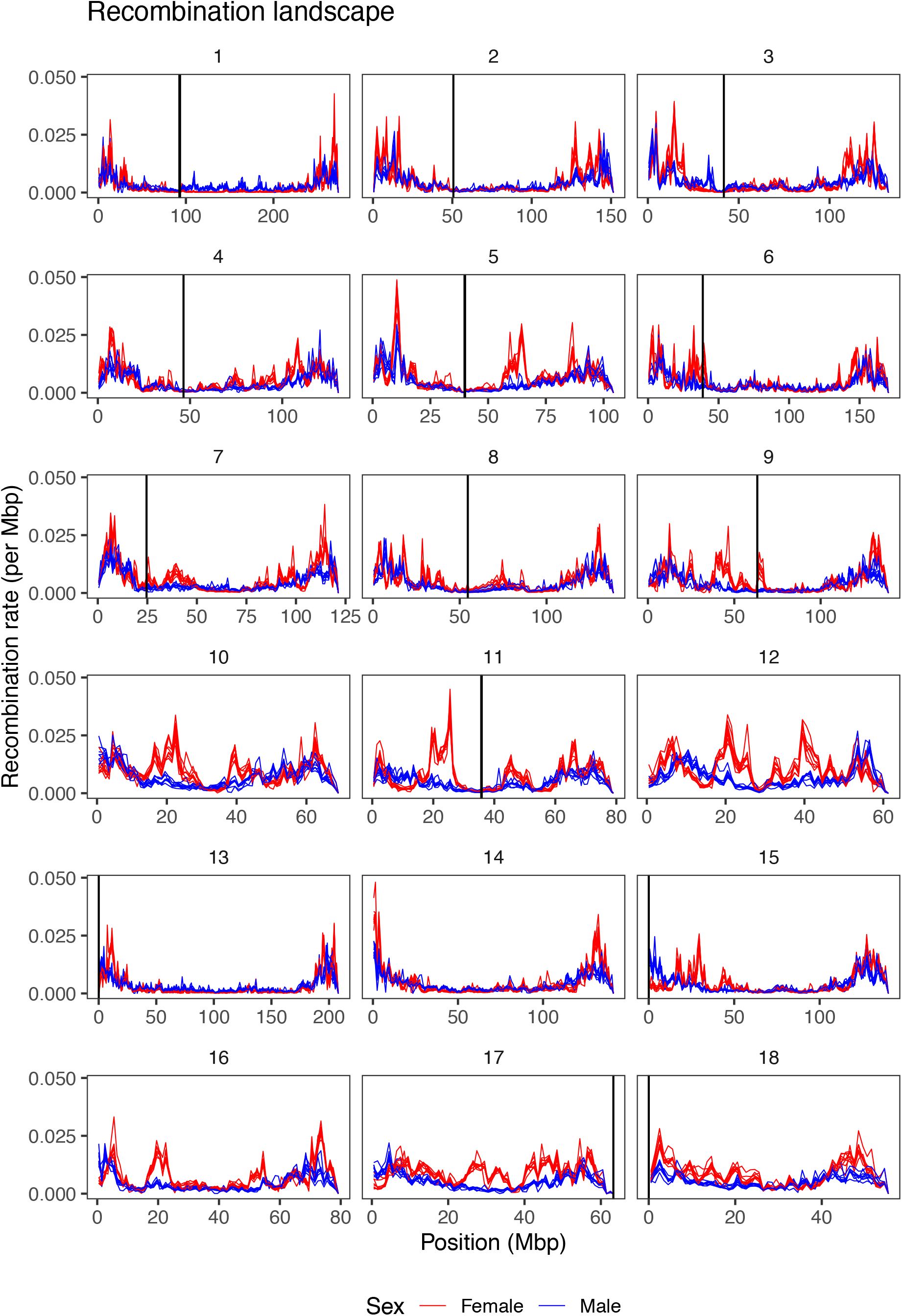
Recombination landscape in the pig. The lines show recombination rate in windows of 1 Mbp along the pig genome (Sscrofa11.1). Red lines show female estimates and blue lines show male estimates. Each line shows one of the nine breeding lines. The black vertical lines are predicted centromere locations in the reference genome, for chromosomes where they are available.

**Figure 3.**
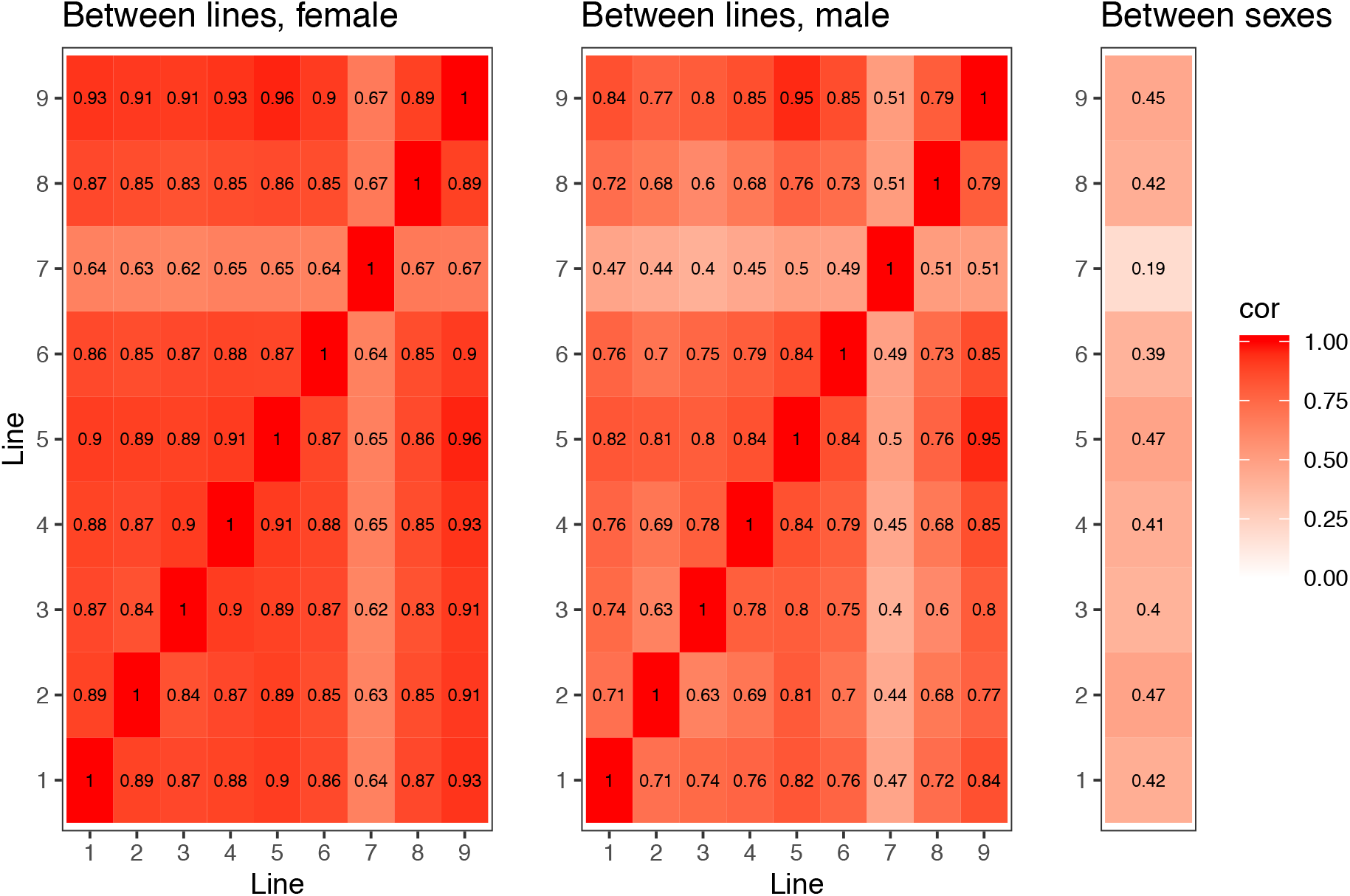
Correlation heatmap of recombination landscapes between lines and sexes. Heatmaps show pairwise correlations between lines of the estimated recombination rates at each marker interval, within each sex, and the correlation between sexes within each line.

### Correlation between genomic features and recombination rate

Local recombination rate had moderate to low correlation (absolute correlation coefficients less than 0.33) with GC content, repeats and particular sequence motifs. Figure 4 shows the correlations between recombination rate and genomic features in 1 Mbp windows, separated by sex. There were positive correlations with GC content, and negative correlation with sequence repeats when all repeat classes were combined. The correlation between recombination rate and different types of repeats was variable. Recombination rate was only weakly correlated with counts of the PRDM9 consensus motif CCNCCNTNNCCNC, but moderately correlated with counts of the CCCCACCCC motif, previously found to be enriched in high recombination regions in the pig genome [1].

**Figure 4.**
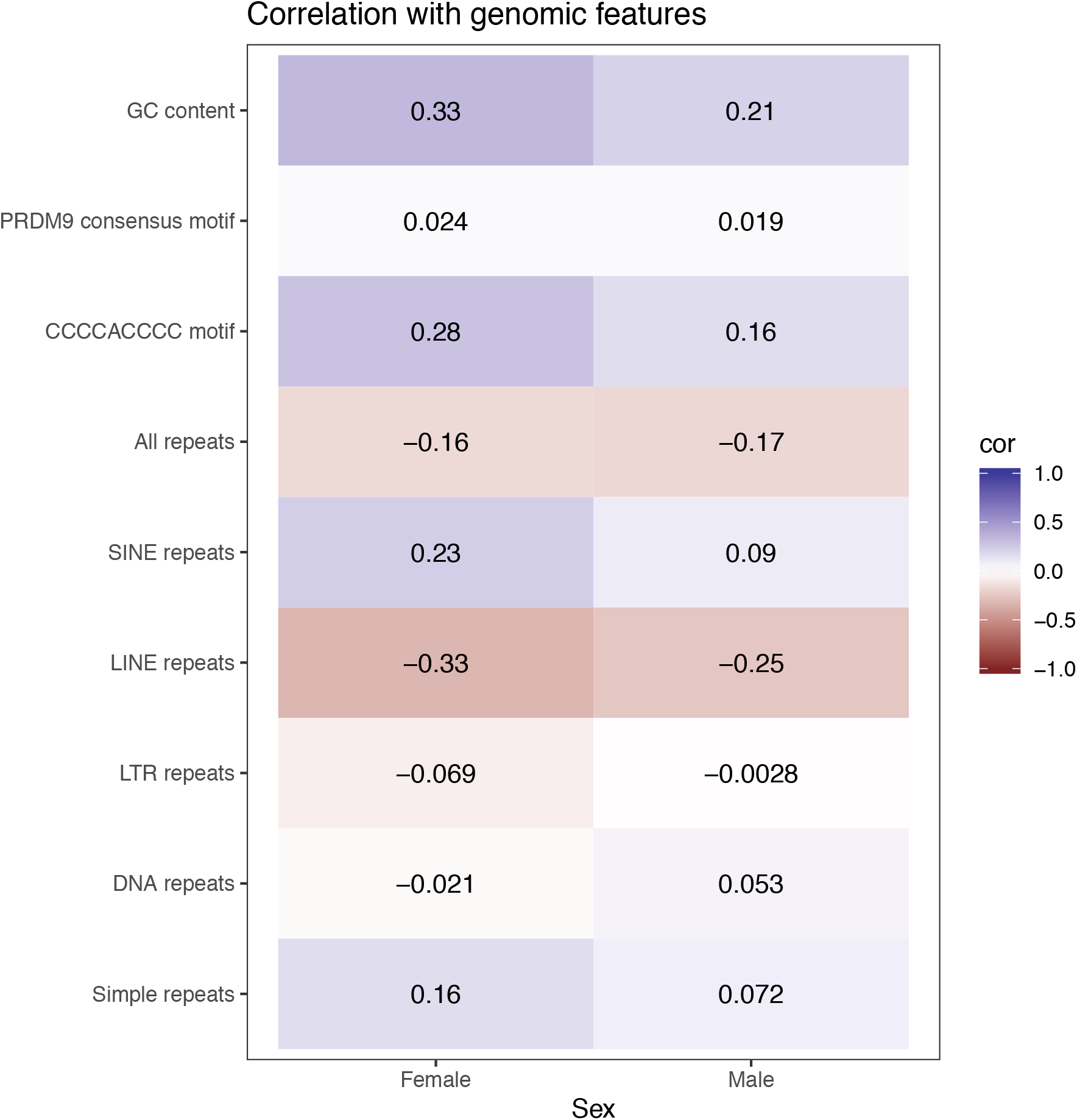
Heatmap of correlation between genome features and recombination rate in windows of 1 Mbp. The heatmap shows correlation between recombination rate sequence features within 2272 windows of the autosomal part of the pig genome (Sscrofa11.1).

### Heritability of recombination rate

Genome-wide recombination rate had low but nonzero heritability (h^2^ on average 0.07 for females and 0.05 for males). Figure 5 shows the heritability and ratio of permanent environmental variance, broken down by sex and line. There was little evidence of differences in heritability between lines. The open circles in Figure 5 show genomic heritability estimates from genome-wide association analyses. The genomic heritabilities suggest that the SNP chip captured most (on average 83%) of the additive genetic variance in recombination.

**Figure 5.**
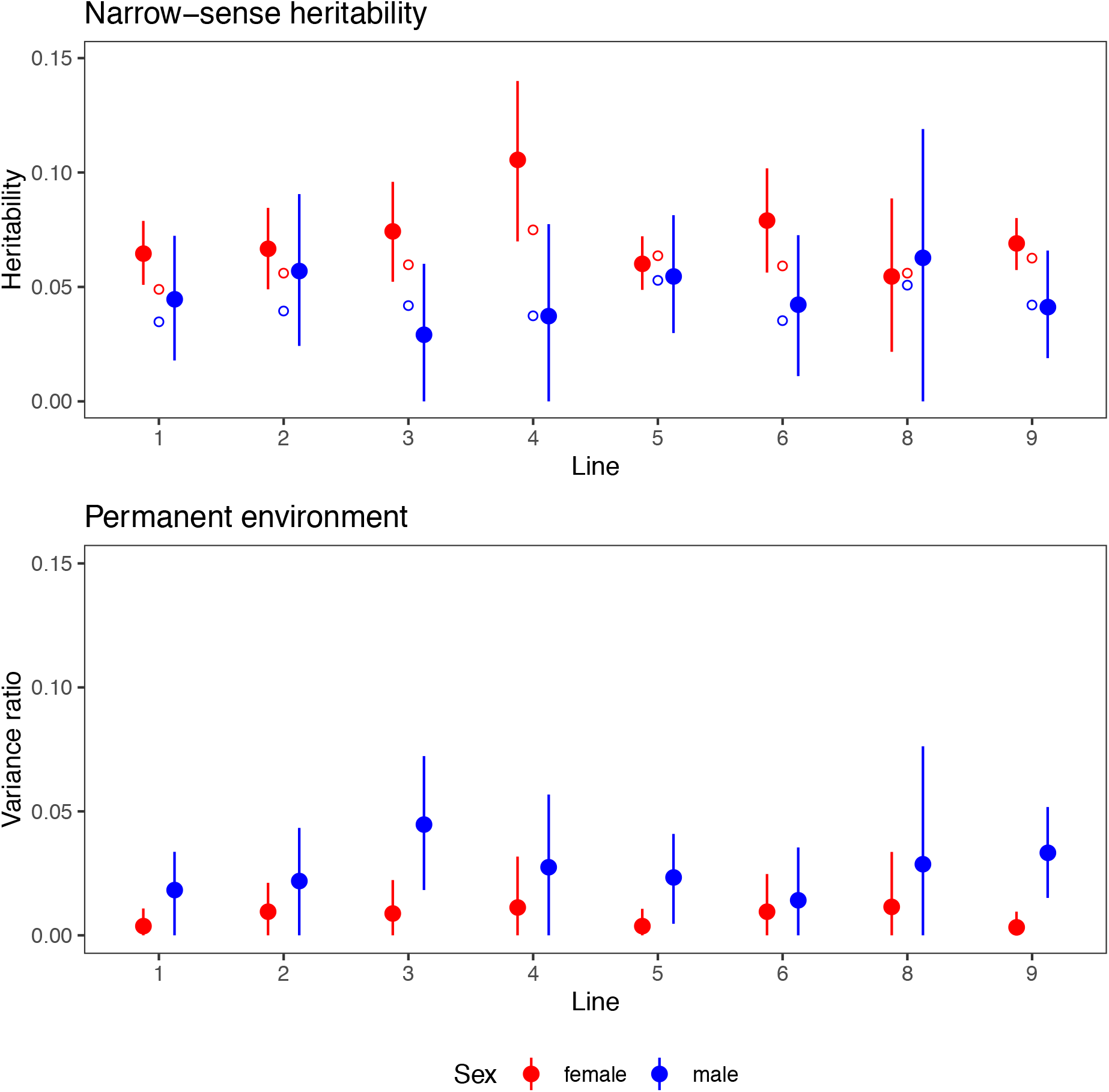
Heritability of average recombination. The dots show estimates of narrow-sense heritability and the permanent environmental effect for average genome-wide recombination estimated with an animal model. The lines show 95% credible intervals. Red dots and lines show female estimates, while blue dots and lines show male estimates. Open circles show the genomic heritability estimated from genome-wide association. Because of the low number of dams and sires, we excluded the smallest line (line 7) from the analysis.

### Genome-wide association of recombination rate

Genome-wide association revealed three regions of the genome containing markers associated with genome-wide recombination rate. Figure 6 shows the results of genome-wide association scans within each line, broken down by sex. Table 3 shows the location of the most significant marker for each region with variance explained and allele frequency. There was a region associated with female recombination rate at the start of chromosome 8 in six of the lines, as well as a region on chromosome 17 in line 1, and one on chromosome 1 in line 6. The chromosome 8 region was also associated with male recombination rate in two lines. Figure 7 shows a zoomed-in view of each of these regions, with the location of known candidate genes involved in recombination.

**Figure 6.**
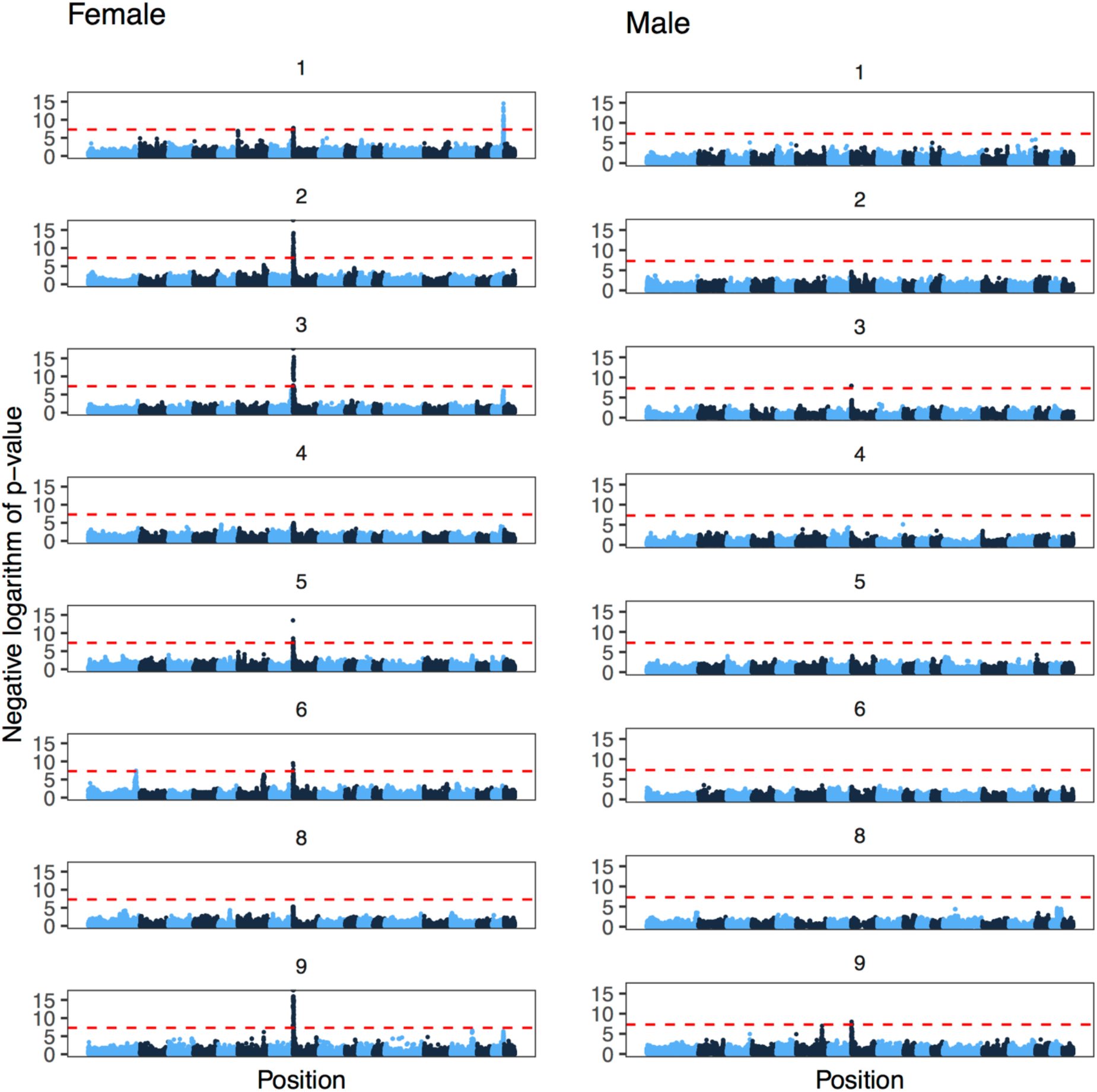
Genome-wide association of average recombination. The subplots are Manhattan plots of the negative logarithm of the p-value of association against genomic position, broken down by line and sex. Alternating colours correspond to chromosomes 1-18. Because of the low number of dams and sires, we excluded the smallest line (line 7) from the analysis. The dashed red line shows a conventional genome-wide significance threshold of 5 · 10^−8^.

**Table 3.** Genome-wide association study hits for average recombination, with position of the lead SNP, additive genetic variance explained by the locus, and allele frequency of the allele associated with higher recombination rate.

| Chromosome | Sex | Line | Lead SNP position | Genetic variance explained | Allele frequency |
| --- | --- | --- | --- | --- | --- |
| 1 | female | 6 | 252,547,401 | 0.10 | 0.57 |
| 8 | female | 1 | 2,253,270 | 0.08 | 0.90 |
| 8 | female | 2 | 75,256 | 0.60 | 0.53 |
| 8 | female | 3 | 226,298 | 0.41 | 0.70 |
| 8 | male | 3 | 226,298 | 0.44 | 0.74 |
| 8 | female | 5 | 259,617 | 0.07 | 0.27 |
| 8 | female | 6 | 259,617 | 0.12 | 0.74 |
| 8 | female | 9 | 75,256 | 0.14 | 0.12 |
| 8 | male | 9 | 1,283,621 | 0.22 | 0.41 |
| 17 | female | 1 | 59,968,884 | 0.16 | 0.78 |

**Figure 7.**
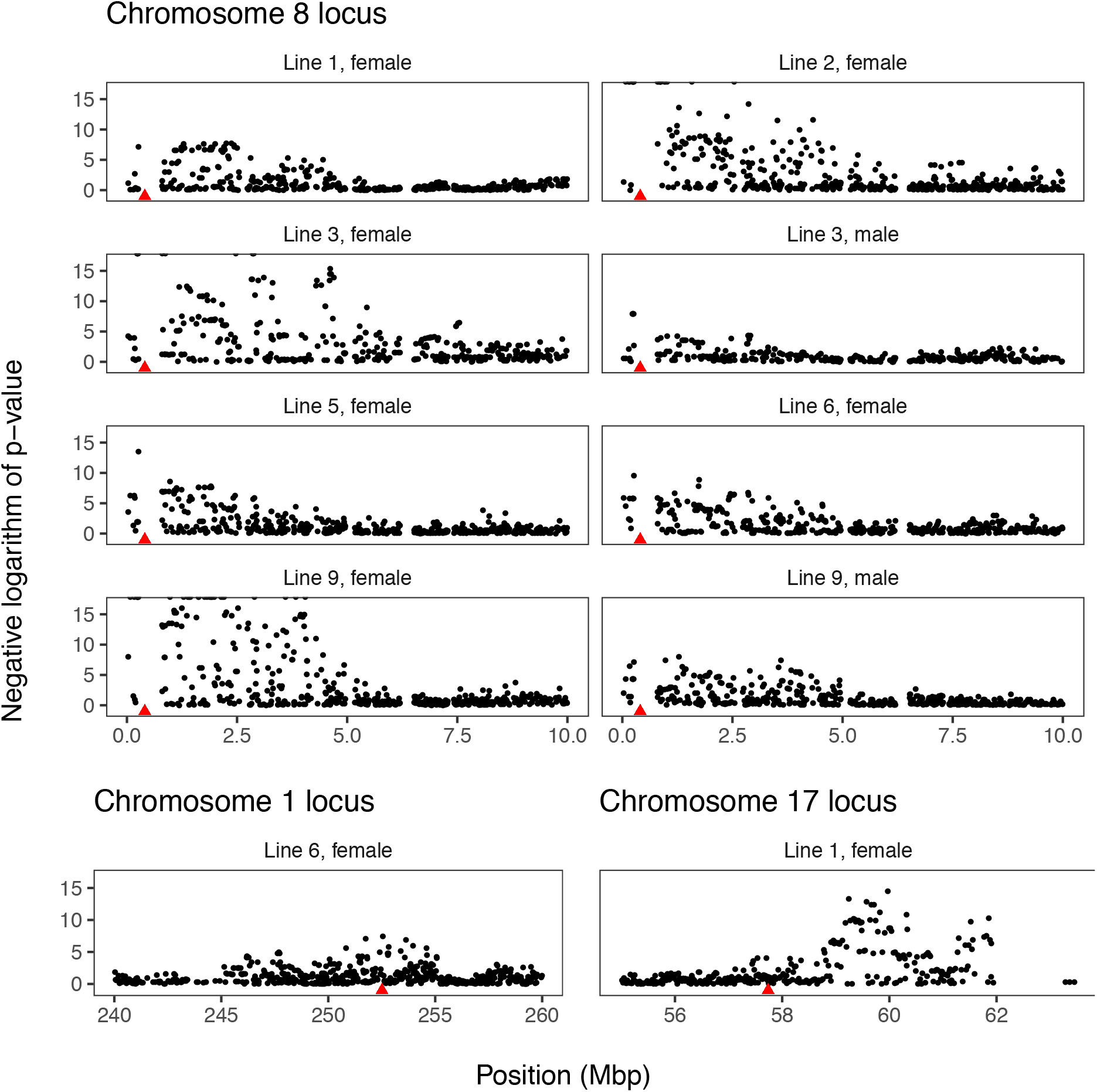
Regions associated with recombination rate and location of recombination-associated candidate genes. The subplots are Manhattan plots of the negative logarithm of the p-value of association against genomic position, zoomed in to show the region around the significant markers. The red triangles show location of *RNF212* on chromosome 8, *SLOC1* on chromosome 1, and *SPO11* on chromosome 17.

### Algorithm performance on synthetic data

We tested the accuracy of the estimated recombination by analysing a synthetic dataset. Figure 8 shows the simulated and estimated map length, recombination landscape, and a scatterplot of simulated and estimated numbers of recombinations per individual. Our method slightly overestimated recombination rate when there was variable recombination along the chromosome. Because of uncertainty in the location of recombinations, the estimated recombination landscape did not track per-marker recombination rate variation very well (r = 0.59), but better captured the smoothed recombination landscape using a window of 50 markers (r = 0.86). The accuracy of individual-level estimates of recombination was higher for dams (r = 0.72) than for sires (r = 0.55).

**Figure 8.**
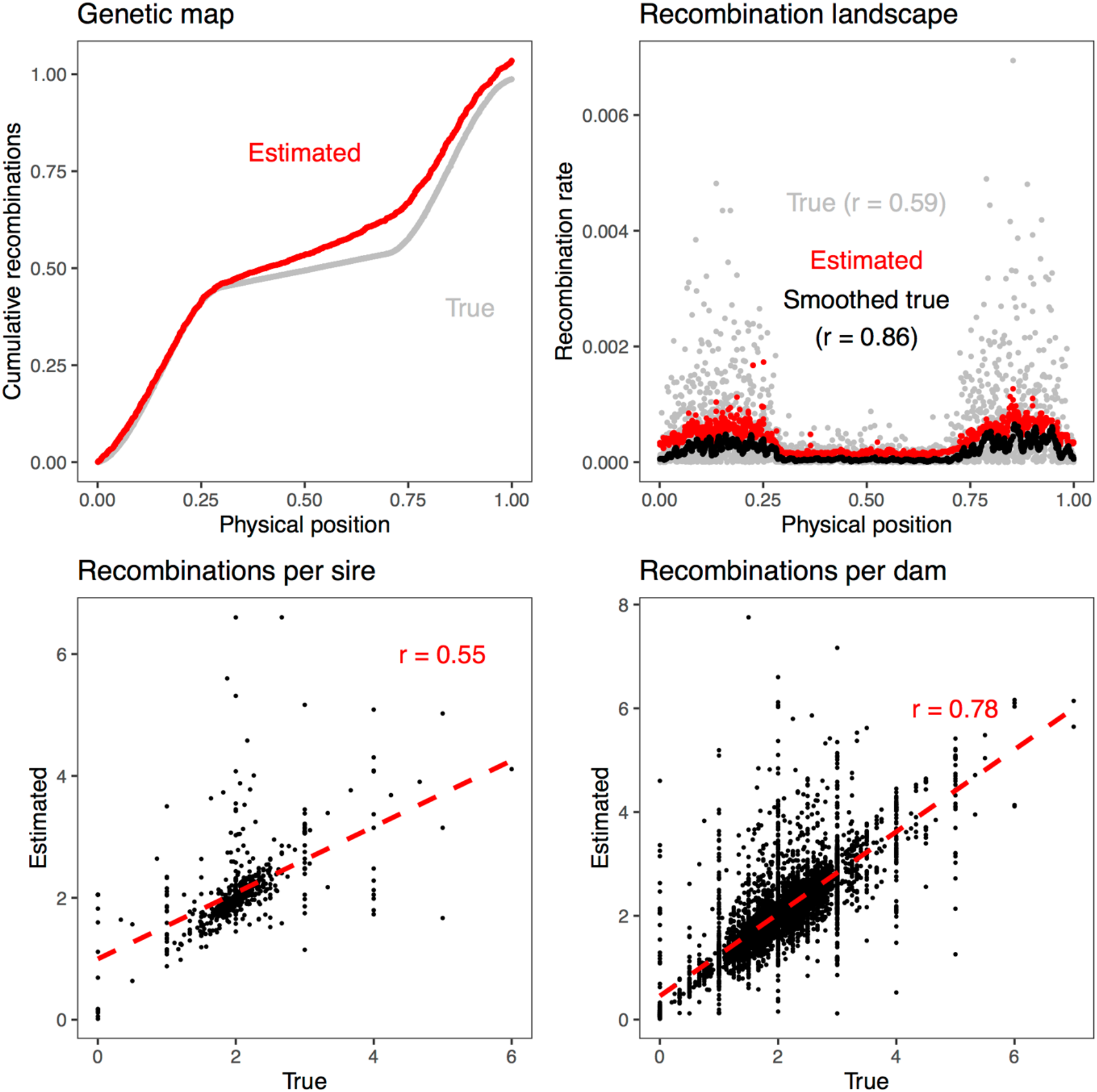
Recombination rate estimation on simulated data. Cumulative number of recombinations, recombination landscape along the simulated chromosome and the correlation between true and estimated numbers of recombination in sires and dams. The smoothed values are rolling averages of 50 markers. The red dashed line is the regression line between true and estimated values.

## Discussion

In this paper, we estimated recombination rate variation within the genome and between individuals in the pig using multiocus iterative peeling in nine genotyped pedigrees.

In this section, we discuss three main results:

1. We confirm the known features of the pig recombination landscape, but not the previously described correlation with the PRDM9 consensus motif.
2. We show that recombination rate in the pig is genetically variable and associated with alleles at the *RNF212* gene.
3. Multilocus iterative peeling is a compelling method for estimating recombination landscapes from large genotyped pedigrees, but tends to overestimate genetic map length.

### Features of the pig recombination landscape

Our results recover known features of recombination in the pig, including the relative chromosome lengths, and the marked sexual dimorphism. There are two notable exceptions, where our estimates disagreed with previous results: we estimate overall longer genetic lengths of chromosomes, and the correlations between recombination rate, density of the PRDM9 consensus binding motif, and the density of some repeat classes are different than estimated previously.

We estimated longer genetic maps than previous estimates for the pig. The total genetic map lengths ranged from 18.5 to 21.7 Morgan for males and 22.3 to 25.9 Morgan for females. In comparison, [1] found sex-specific map lengths of 17.8 and 17.5 Morgan for males, and 22.4 and 25.5 Morgan for females. This may be due to overestimation (see below), but also a higher marker density and more complete use of the pedigree allowing us to detect more recombinations.

The correlation between recombination and density of the PRDM9 consensus binding motif, was lower than previous estimates. Because the PRDM9 protein determines the locations of a subset of recombination hotspots, a positive correlation was expected. We detected only a weak positive correlation with PRDM9 consensus motif density and recombination, which suggests that we lack the genomic resolution to detect variation at this scale. The recombination rate landscape is the outcome of processes operating at a much smaller scale, with hotspots of a few kilobasepairs (as estimated by population sequencing [3] or by high-density gamete genotyping [33]). There is more subtle local variation in recombination rate that we cannot detect.

The associations between recombination and transposable element density were mixed, and different for different types of transposable elements. The overall correlation between recombination rate and repeats was negative, in line with estimates from other species [34]. The negative correlation with LINEs was stronger than previously reported and the positive correlation with simple repeats was weaker. One reason for these differences might be that we used the more complete Sscrofa11.1 reference genome [26], which likely better resolves the repeat landscape of the pig genome than the previous version.

### Genetic variation in genome-wide recombination rate

Our results from the pig agree with the general picture of recombination rate variation in vertebrates. The chromosome 8 locus is homologous to regions identified in humans [35–37], cattle [7, 8, 10], sheep [12, 13], and chickens [14]. It contains the *RNF212* gene, a paralog of which is also associated with recombination in deer [11]. The RNF212 protein binds to recombination complexes, and is essential for crossover formation [38].

While *RNF212* is an obvious candidate gene, it is harder to find candidates for the other two regions. We searched for the locations of candidate regions from other vertebrates, and rapidly evolving recombination genes in mammals [39]. The chromosome 1 locus overlaps *SHOC1,* one of the rapidly evolving recombination genes in mammals [39]. The closest candidate recombination gene from the chromosome 17 locus is *SPO11,* associated with recombination in chickens [14]. However, it is about two megabasepairs away from the most significant marker.

There are differences in recombination rate between lines, which may be due to fixed genetic differences. Given that livestock populations have relatively small effective population sizes, and assuming that recombination rate variation has a rather simple genetic architecture, line differences in recombination rate might very well be due to genetic differences that have fixed by chance. At the same times, all the lines showed evidence of comparable genetic variation in recombination rate, and there was evidence that the major locus on chromosome 8 segregates in most lines.

A higher recombination rate could be beneficial for breeding, because it would reduce linkage disequilibrium between causative variants and release genetic variance. Simulations suggest that substantial increases in genome-wide recombination rate could increase genetic gain [40]. We can approximate how much breeding could increase recombination rate based on our results.

First, we can use the Breeder’s equation to predict the response to selection, treating genome-wide recombination as a quantitative trait. The response is the heritability multiplied by the selection differential *S*, which is the difference between population mean *μ* and mean of the selected individuals *μ*_*selected*_.

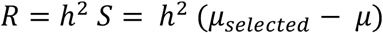

Using distribution of genome-wide recombination rates from the males of the largest line, the mean were 0.904 cM/Mbp. If we were to select the 10%, 20% or 30% highest recombination individuals, the mean of the selected individuals would be 1.22 cM/Mbp, 1.15 cM/Mbp, and 1.11 cM/Mbp respectively. Assuming a heritability of 0.05, comparable to our estimated genomic heritability, this would result in responses of:

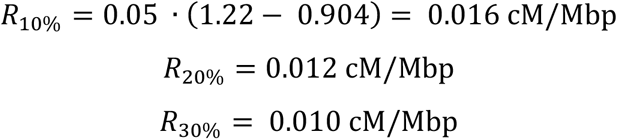

Relative to the average recombination rate, that would mean increases of 1.7%, 1.3% and 1.1%, respectively.

Second, we concentrate on the major locus on chromosome 8 that we detected in most of the lines, and approximate the increase in recombination rate that could be achieved if this locus was fixed for the high recombination allele. Again, using estimates from the largest line, the additive effect *a* of the chromosome 8 locus was estimated to be 0.0271 cM/Mbp (averaging the male and female estimates), and the frequency *f* of the high recombination allele was 0.332 (weighted average of males and females). The increase in the mean of the population by fixing the chromosome 8 locus would be:

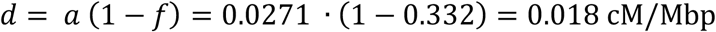

That is, it would increase genome-wide recombination rate by about 2%.

Compared to the simulation results of [40], which suggest that a doubling or more of genome-wide recombination rate would lead to substantial genetic gains, these results suggest that breeding for higher genome-wide recombination rate is not a practical alternative for improving genetic gain. There may be other potential avenues, such as introducing targeted recombinations in favourable locations [41] by biotechnological means.

### Recombination rate inference by multilocus peeling

Throughout this paper we have used multilocus iterative peeling to estimate recombination rate. In our simulation study, we found that multilocus iterative peeling could estimate the number of recombinations per individual with an accuracy of 0.7 for dams and 0.5 for sires, and the average recombination landscape along a chromosome. This is consistent with our analysis of the pig genome, where we confirm previously known features of the pig recombination landscape. However, the simulation results also show that we overestimated the total genetic map length, consistent with our comparisons between the estimate recombination rate and previously published estimates [1].

Multilocus iterative peeling presents a compelling technique for estimating recombination rate in large pedigree populations: it scales well to massive livestock pedigrees (more than 150,000 individuals), does not require pre-phasing of the data, and handles individuals genotyped on range of platforms without requiring non-overlapping variants to be imputed beforehand.

The primary downside is that multilocus iterative peeling requires multiple generations of genotyped individuals to be available to accurately phase, impute, and estimate the recombination rate. Although this information may be available in pig or chicken breeding programmes [23, 42], and some wild populations [12] it may not be available in all populations. In addition to this the overestimation of genetic map length suggests that the exact genetic map lengths and counts of recombination for a specific individual may not be accurate, but it is able to recover broad patterns in recombination between chromosomes and between individuals.

## Conclusion

In this paper we analyse 150,000 individuals from nine pig pedigrees. We find that we are able to recover broad-scale patterns in the total genetic map length, recombination landscape, and sex differences in recombination rates. In addition to this, we found that recombination rate had low, but non-zero heritability, and a genome-wide association study detected three regions associated with recombination rate. This paper highlights the ability to use large scale pedigree and genomic data, as is routinely collected in many closely managed populations to infer and understand recombination and recombination rate variation.

## Declarations

### Ethics approval and consent to participate

The samples used in this study were derived from the routine breeding activities of PIC.

### Consent for publication

Not applicable.

### Availability of data and materials

The datasets generated and analysed in this study are derived from the PIC breeding programme and not publicly available.

### Competing interests

The authors declare that they have no competing interests.

### Funding

The authors acknowledge the financial support from the BBSRC ISPG to The Roslin Institute BBS/E/D/30002275, from Grant Nos. BB/N015339/1, BB/L020467/1, BB/M009254/1, from Genus PLC, Innovate UK, and from the Swedish Research Council Formas Dnr 2016-01386.

### Author’s contributions

JMH, MJ, AW and GG conceived the study. MJ, AW, RRF and CC analysed data. WH and DdK helped interpret the results. MJ, AW and JMH wrote the paper. All authors read and approved the final manuscript.

## Acknowledgements

This work has made use of the resources provided by the Edinburgh Compute and Data Facility (ECDF) (http://www.ecdf.ed.ac.uk).

